# Estimation of neuronal tuning for word meaning from passively recorded naturalistic speech

**DOI:** 10.64898/2026.06.23.733980

**Authors:** Taha Ismail, Ana G. Chavez, Xinyuan Yan, Hanlin Zhu, Melissa Franch, James Belanger, Saipravallika Chamarthi, Katherine E. Kabotyanski, Kalman A. Katlowitz, Assia Chericoni, Elizabeth A. Mickiewicz, Timon Merk, Yewen Zhou, Nikhil Shivakumar, Paul J. Steffan, Rahul Hingorani, Mattson Ogg, Han Yi, Tomasz Fraczek, Eleonora Bartoli, Jay A. Hennig, Sameer A. Sheth, Nicole R. Provenza, Benjamin Y. Hayden

**Affiliations:** Department of Neurosurgery, Baylor College of Medicine; Department of Neuroscience, Baylor College of Medicine; Department of Psychiatry and Behavioral Sciences, Baylor College of Medicine; Department of Electrical and Computer Engineering, Rice University; Neuroengineering Initiative, Rice University; Department of Bioengineering, Rice University; Research and Exploratory Development Department, Johns Hopkins University Applied Physics Laboratory

**Author notes:** Co-equal authorship (Sameer A. Sheth, Nicole R. Provenza, Benjamin Y. Hayden).

## Abstract

The ability to derive neural-level language coding models holds great scientific and clinical potential. Current approaches are limited by the scale and ethological validity of input data; applications requiring large, rare, or naturalistic samples in particular would benefit from the ability to infer neural coding from incidental everyday speech. Here we present a novel pipeline designed to leverage spontaneous and incidental naturalistic speech. This pipeline performs transcription, segmentation, and video-assisted diarization, as well as alignment and spike detection of neural data. We apply this pipeline to a dataset derived from 21 patients (6+ days each, over 800 hours and 5 million words total). We benchmark both encoding and decoding models against extensive and rare ground-truth control datasets consisting of human-curated word-level temporal alignment and manually sorted spikes. We further validate our approach by quantifying representational drift, effect of dataset size, and differences between six brain areas. Together, these findings demonstrate that incidental natural speech is sufficiently processed in the brain to enable the estimation neural-level embeddings.

## INTRODUCTION

As we listen to speech, our brains continuously transform that input into structured neural representations. By combining measurements of neural responses with embeddings derived from large-language models, scientists can quantify the association between neural responses and the language that evokes them. This approach works with a range of neural measures, including fMRI, MEG, EEG, and intracranial EEG (Hasson et al., 2020; Huth et al., 2016; Caucheteux & King, 2022; Hasson et al., 2020; Franch et al., 2025; Jamali et al., 2024; Toneva et al., 2020; Wehbe et al., 2014; Zeng & Gallant, 2025). While each of these measures have their value, neuron-level recordings track both the high temporal resolution of speech and give the high spatial resolution of neural computation (Fried et al., 2014; Rutishauser et al., 2015; Franch et al., 2025; Jamali et al., 2024; Quian Quiroga et al., 2005; Katlowitz et al., 2025; Yan et al., 2025). As a result, they are often used in brain-computer interfaces (BCIs, Card et al., 2024; Willett et al., 2023; Moses et al., 2021), as well as in basic science studies focused on mechanism (e.g., Zhu et al., 2026).

Neuron-level language representations can be formalized using *encoding models*, which relate patterns of brain activity to language model-derived embeddings (Huth et al., 2016; Mitchell et al., 2008; Wehbe et al., 2014; Goldstein et al., 2022; Franch et al., 2025; Jamali et al., 2024; Caucheteux et al., 2021; Goldstein et al., 2025). *Decoding models* provide a complementary perspective by asking whether neural data can classify linguistic stimuli (Naselaris et al., 2011; Huth et al., 2016; Makin et al., 2020; Willett et al., 2021). Both have potential value for both clinical and basic science; moreover, their joint application potentially allows for both improved interpretability and enhanced understanding (Kriegeskorte & Douglas, 2019; Holdgraf et al., 2017).

Most such studies rely on brief, highly controlled laboratory paradigms with laborious manual curation of speech (transcription, segmentation, diarization) and manual processing of neural data. As a result, it remains unknown whether neural-level robust semantic models can be recovered from long-duration, naturalistic speech encountered in everyday life, and, if so, how accurate they will be (Sonkusare et al., 2019; Nastase et al., 2020; Russell et al., 2024; Matusz et al., 2019; Hamilton & Huth, 2018). Modeling neural responses from incidental continuously recorded speech introduces several challenges (Evanson et al., 2025; Sonkusare et al., 2019; Goldstein et al., 2025). Everyday speech contains frequent dysfluencies, variable articulation, speaker overlap, and shifting acoustic contexts, all of which complicate precise temporal alignment between language and neural activity (Shriberg, 2001; Giraud & Poeppel, 2012; Mesgarani & Chang., 2012; Ding & Simon., 2014). These properties make accurate transcription, diarization, and segmentation difficult. Moreover, the long durations of natural conversation preclude gold-standard manual spike sorting across long recordings (Jun et al., 2017; Pachitariu et al., 2016). Although emerging automated pipelines for speech processing and spike detection now make approximate spike sorting in such datasets tractable, these tools remain imperfect and can introduce additional sources of noise and bias (Bain et al., 2023; Bredin, 2023; Kuchaiev et al., 2019; Pacheco et al., 2025; Pachitariu et al., 2024; Magland et al., 2020; Garcia et al., 2022; De Preter et al., 2024). One of the biggest barriers to evaluating these models is the lack of high quality ground-truthed datasets large enough to evaluate automated pipelines. Despite these barriers, we hypothesize that the brain processes natural speech well enough that automated pipelines can reliably be deployed to recover valid speech models.

In any case, naturalistic speech offers unique scientific and practical advantages. Daily life provides an exceptionally rich and diverse linguistic input—often tens of thousands of words per day—far exceeding what can be presented in conventional experiments (Evanson et al., 2025; Goldstein et al., 2025; Nastase et al., 2021; Sonkusare et al, 2019). This scale is critical for studying rare linguistic events, context-dependent meaning, and phenomena such as dysfluencies that can be difficult to elicit under controlled conditions. In addition, continuously updated models derived from everyday speech are increasingly relevant for clinical technologies, including speech prosthetics and adaptive deep brain stimulation, which require robust neural–linguistic mappings outside the laboratory (Card et al., 2024; Herron et al., 2017; Pina-Fuentes et al., 2020). Here we describe a novel pipeline deployed to analyze all incidental speech from within the EMU at a single hospital for 21 patients over 800 hours. Critically, we leverage a ground-truth dataset derived from real-world conversations that is manually aligned and diarized, along with corresponding manual spike sorting.

## RESULTS

We recorded speech and brain activity over a period of 6 to 11 days for 21 adult patients with medically refractory epilepsy undergoing intracranial seizure monitoring. We continuously recorded speech occurring during the entirety of each patient’s stay (average: 6.72 +/- 0.52 days) in our epilepsy monitoring unit (EMU, **Methods**). Two studio microphones, one directly above the patient’s bed and another to the side, recorded 24/7 speech occurring within the patient room. Additionally, eight cameras located throughout the room, capturing various angles of the patient bed, recorded patient activity during their stay (**Figure 1A**). The microphones and cameras recorded many conversations with friends, family, clinical and research staff, as well as speech played from the television (**Figure 1B**). Intracranial EEG electrodes with microwires for collecting single-neuron spiking data (median: 56 contacts per patient, ranging from 16-96 contacts in various brain regions) recorded single and multi-unit spiking activity at 30 kHz for the entire patient stay, allowing us to sync microwire unit data at high temporal resolution.

**Figure 1:**
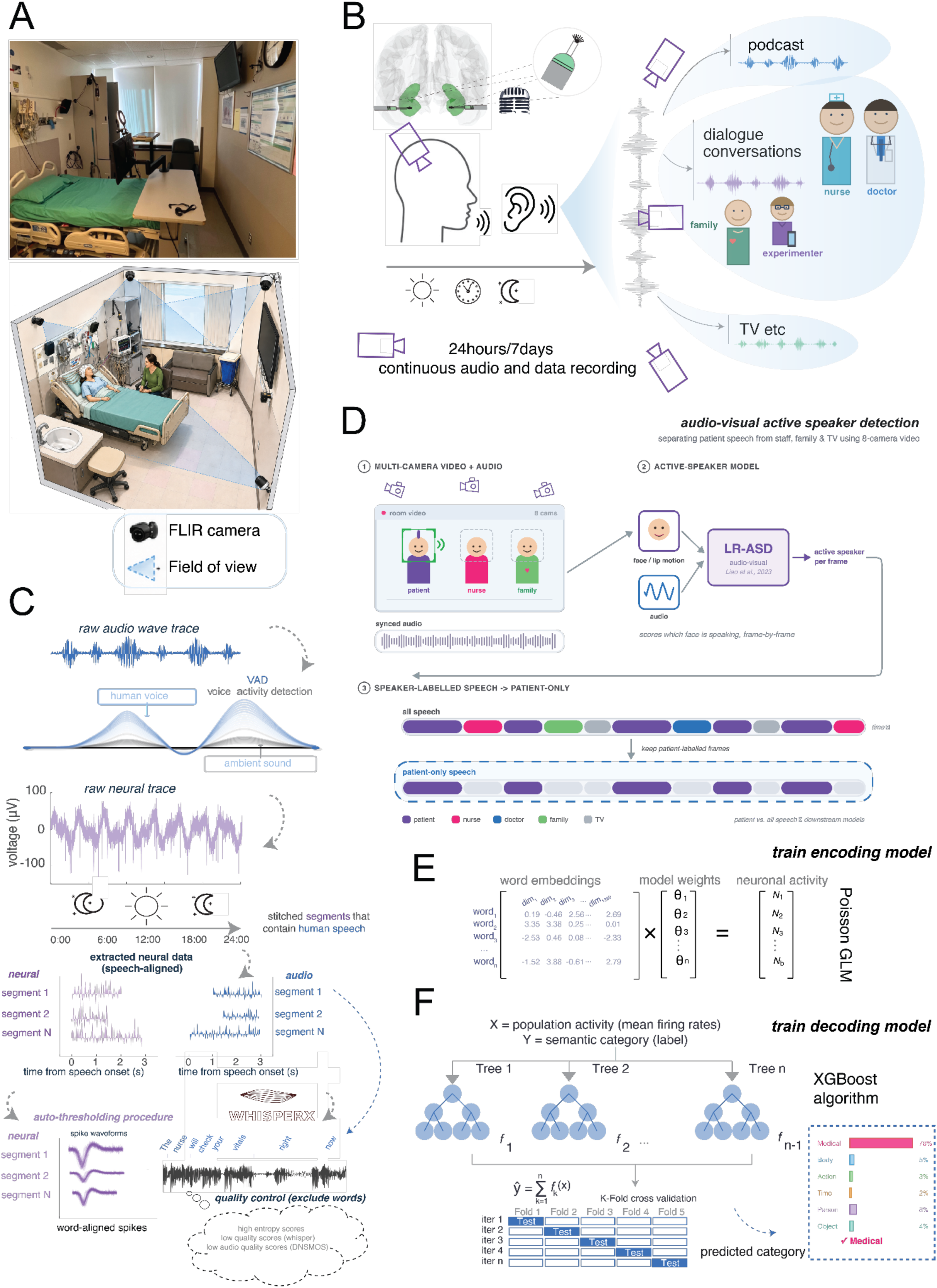
Neural-behavioral Sync and Modeling Pipeline. **A.** Picture and diagram showing the Epilepsy Monitoring Unit (EMU) where patients are kept for intracranial seizure monitoring, with icons displaying the audio and video recording modalities present in the room and for research use. **B.** Synchronization of audio and neural signal courtesy of our Blackrock Neurotech recording system that maintains a synchronized stream of both, sampled at 30 kHz **C.** Automated speech extraction and autothresholding pipeline. **D.** Methodology for extracting patient speaker scores from the audio-video stream and filtering patient-only speech segments **E.** Input structure of our Poisson GLM used for semantic and emotion encoding **F.** Input and model structure used for XGBoost semantic category decoding.

All speech data were processed through a voice activity detection model (Silero Team, 2024) to find passages of audio data that contained identifiable human speech. Speech was then segmented into sentences and words using WhisperX (Bain et al., 2023). Multiple scoring metrics, including CTC log-likelihood, WhisperX quality score, spectral entropy, and DNSMOS quality scores were calculated for each sentence (**Methods**). Sentences at low-quality extrema (outside 0.75 * IQR) of their relative distributions were removed from the dataset (**Figure 1C**). We used a standard autothresholding algorithm (Rossant et al., 2016; Rutishauser et al., 2014; Hammad et a., 2013) to capture multi-unit spiking activity that could be aligned with word timestamps (**Figure 1C**, **Methods**). For convenience, we refer to each of these autolabelled sets as *units*.

To identify segments of patient speech, we used a multimodal active speaker detection (ASD) model (Liao et al., 2023). Audio-only diarization methods make frequent errors in naturalistic recording conditions, such as overlapping speakers and background noise, which led to us turning to multimodal methods (Cheng et al., 2023; Park et al., 2022). Specifically, we combined video with audio features to detect facial movements associated with speech to identify video frames in which the patient is likely to be speaking (**Figure 1D, Methods**). We used the speaker labels from this model to compare patient-only versus all speech in our downstream semantic coding models.

Our **encoding models** involved regressing GPT-2 embeddings against spike counts for individual units (cf. Franch et al., 2026; Katlowitz et al., 2025; Yan et al., 2025). The model was implemented as a Poisson GLM with an L2 regularizer and a duration term included as an offset variable (**Figure 1E**). Models were fitted independently across neurons in bulk matrices in PyTorch (**Methods**). GPT-2 embeddings were calculated by taking the hidden state of the last layer of GPT-2-large from an inference pass of the current word, with the prior 200 words in the continuous speech stream taken as context (**Methods**). The final hidden layer was selected due to it being the natural contextual representation of a word as it is directly used for next-token prediction in the transformer architecture (Vaswani et al., 2017). Moreover, prior studies find qualitatively robust performance for neural encoders in the final layer of GPT-2 as well as earlier layers (Katlowitz et al., 2026; French et al., 2025; Yan et al., 2026)

Our **decoding models** involved training an XGBoost classifier with population firing rate activity (defined as the average neural firing rate across a word interval for a set of neurons) to predict broad semantic category labels, which included classes such as “relationships”, “numerical”, and “medical” (**Figure 1F**). XGBoost was chosen for its skill in learning nonlinear classifier boundaries, which is helpful for neural decoding (Chen et al., 2016). Moreover, XGBoost is frequently used with success for behavioral decoding from invasive neural recordings (Merk et al., 2022; Hirschmann et al., 2022).

### Dataset spanning millions of words with accurate transcription and diarization

We recorded a total of 5,270,665 words both spoken and heard by 21 participants with simultaneous brain recordings. The number of words for each patient ranged from 40,351 to 521,360. Seventeen of the patients had datasets with over 100,000 words. Across all patients, we collected a total of 871.2 hours of speech (spoken and/or heard), ranging between 7.1 and over 87.7 hours of speech per patient (**Figure 2A**). Across the fourteen patients for which we have quality video data available, we collected a total of 156,397 words of patient-specific speech, ranging from 1,644 to 33,972 words per patient. We find the lexicon of spoken words to span a range of expected topics, including a large mode for medical terminology (**Figure 2B**). Additionally, we find that the timing of speech production and listening follows a cyclic pattern, with the number of words spoken peaking in the early afternoon hours and declining significantly late at night (**Figure 2C**).

**Figure 2:**
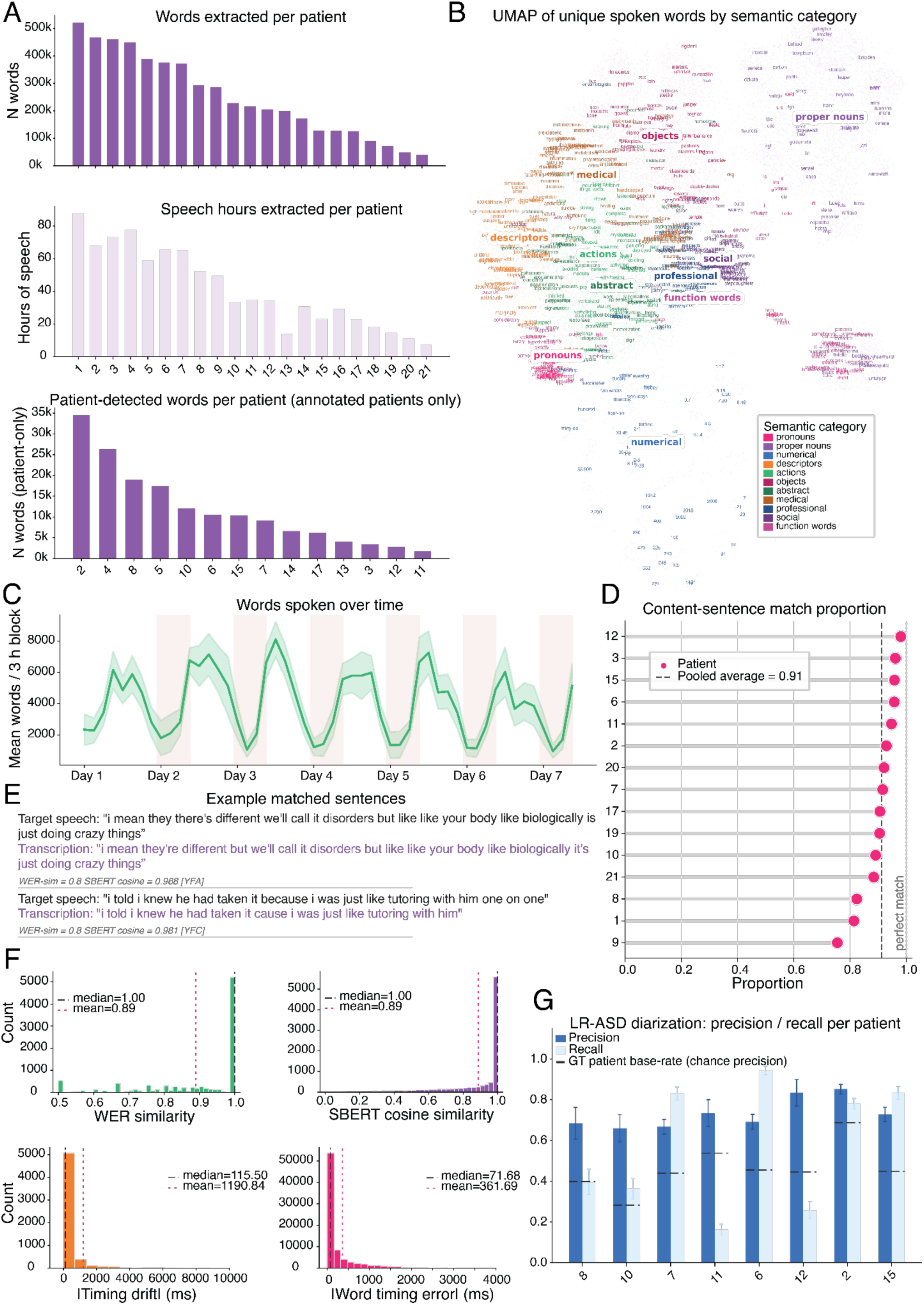
Data quantity and annotation accuracy. **A.** 2D UMAP plot of all words collected from our pipeline, clustered by word2vec embeddings and colored by semantic category. Words generally cluster by their assigned semantic category **B.** Bar plots showing the amount of speech data collected from our pipeline per patient, sorted by number of words. **C.** Line plot showing the average number of words across patients collected per 3 hour block starting the first midnight after patient admission, displaying a daily cyclic pattern. Background color displays the standard error per time block across patients. **D.** Lollipop plot displaying the portion of conversation sentences for which a corresponding match was found in the associated automated transcript. **E.** Example sentences displaying what potential pairs of matched sentences looked like and the types of transcription errors corrected for. **F.** Histograms describing transcription and segmentation errors across matched sentences in terms of WER similarity, SBERT cosine similarity, sentence timing errors, and word timing errors. **G.** Paired barplot summarizing diarization accuracy of annotated conversations compared to patient base speaking rate.

***Ground-truth accuracy:*** We first quantified the accuracy of the WhisperX transcription and video-based diarization for our specific EMU-based context. For a subset of fifteen patients, we used sections of data in which study staff had planned conversations with the patient (∼45 minutes/patient). We then performed precise manual temporal alignment and speaker identification (i.e., diarization) of every word spoken and heard (Chavez et al., 2025). We used these data as a ground truth to evaluate the accuracy of the automated transcription and diarization of the same conversation. For transcription accuracy, we evaluated every sentence in our manual transcript with its corresponding match in the automatic transcript. We focused on semantically meaningful sentences, which included sentences with at least two non-filler words (filler words here refers to words like “*um*”, “*hmm*”, and “*haha*”). For each sentence in the manual transcript we searched for its corresponding sentence (if any) in the automated transcript within a 60 second window around the sentence. A match was considered to be the sentence with the lowest word error rate (WER) in the window as long as it was 0.5 or less, which was determined via Levenshtein distances (Levenshtein, 1966, **Methods**). We found that, on average, 90.15% of semantically meaningful sentences were matched in the automated transcript, with the automated transcript capturing the majority of sentences in the conversation for all patients (**Figure 2D**). This correspondence indicates WhisperX is generally successful in capturing instances of speech in the EMU, with infrequent errors resulting from known variables such as overlapping speakers and noisy audio segments (Bain et al., 2023).

***Quantifying semantic precision:*** We calculated four summary statistics for sentence matches in order to assess the extent of transcription errors. The average word error rate similarity score was 0.89, indicating that only 11% of the characters in automated transcripts differed from the true transcript on average. We next calculated the cosine similarity between matched sentence embeddings (derived via the all-mpnet-base-v2 version of the SBERT sentence transformers model; Reimers et al., 2019). We found the mean cosine similarity score to also be 0.89. When looking at non-perfect matches (i.e., WER similarity < 1.0), the average SBERT cosine similar score is greater than the average WER similarity score (p<0.001). As such, even in the cases where minor word-level or syntactical errors are made, semantic meaning tended to be maintained for sentence matches. For example, the sentences “I knew he had taken it *because* I was just like tutoring with him *one on one*” and “I knew he had taken it *cause* I was just like tutoring with him,” had a slightly lower WER similarity score of 0.8 whereas their SBERT cosine similarity is 0.981 (**Figure 2E**).

***Quantifying temporal precision:*** We also calculated the timestamp error of all matched sentences and words within those sentences to quantify temporal accuracy in word segmentation. Notably, the median absolute distance between matched sentences (defined from the midpoints of the sentences) was 115.50 milliseconds, and the median absolute distance between words within each matched sentence was only 71.68 milliseconds. This minimal discrepancy indicates timestamp errors are relatively minor and should not significantly affect estimates of semantic tuning. Overall, we found that transcription and segmentation errors were relatively minor and should not invalidate the possibility of generating semantic coding models from naturalistic speech (**Figure 2F**).

To assess our diarization accuracy, we compared the outputs of LR-ASD to the manually annotated speakers for each sentence. LR-ASD provides a speaker confidence score for every frame of a video, so we labeled a speech segment as coming from the patient if the model labeled the patient as speaking for the majority of frames. Labelling in this way notably struggles in sections with overlapping speakers, similar to automated transcription. When calculating basic classification metrics and comparing them to each patient’s base speaking rate (i.e., the portion of the conversation they spent talking), we find that the video-diarization model is significantly more precise than the base speaking rate for all tested patients for which we had video data available (**Figure 2G**). Moreover, for 50% of patients with video available, LR-ASD is more than 70% accurate when classifying a sentence as belonging to the patient. This result is in line with other groups’ findings that multimodal diarization models significantly outperform audio-only methods (He et al., 2022; Chung et al., 2020). We find that our pipeline struggles with recall for many patients dependent on video quality, meaning that while we can be reasonably confident that a sentence marked as “patient” is patient speech, we cannot be confident that we are necessarily detecting the majority of patient-spoken sentences. As such, for downstream analyses we do not make the assumption that sentences not classified as patient speech necessarily are spoken by other people. Overall, while automated diarization is not as comprehensively accurate as automated transcription, we consider it to be sufficiently reliable for analyses evaluating semantic tuning of patient-specific speech.

### Significant semantic encoding in the continuous speech dataset

We first aimed to understand whether our automated pipeline allows for the development of meaningful semantic encoding models. We analyzed recovered autothresholded unit data from all available brain structures: (1) hippocampus (HPC, 496 units), (2) amygdala (AMY, 88 units), (3) orbitofrontal cortex (OFC, 104 units), (4) thalamus (THAL, centromedian nucleus, 64 units), (5) anterior cingulate cortex (ACC, 240 units), and (6) posterior cingulate cortex (PCC, 24 units). We first examined all autothresholded units together. We collected between 16 to 64 units per patient, with the most units coming from the hippocampus and anterior cingulate and the least coming from the thalamus and posterior cingulate.

We evaluated *<u>encoder model performance</u>* using the pseudo R^2^ score, which defines the relative performance gain over a null model that predicts spike counts from the average firing rate and duration of each word (**Methods**). Pseudo R^2^ was chosen for its interpretability and inherent benchmarking: a score equal to zero indicates performance that is equal to our specified null model. Negative values indicate performance that was worse than the null model. Moreover, Pseudo R^2^ scores with mean firing rate nulls are commonly used to evaluate GLM performance in neural coding analyses (Kraus et al., 2015; Benjamin et al., 2018; Mimica et al., 2023). Our automated pipeline produced robust encoding models, as seen from average pseudo R^2^ scores above 0 (0.004 ± 0.0003 to 0.12 ± 0.025) for all patients (**Figure 3A**). Our response range is typical for neural encoding models (Kraus et al., 2015; Benjamin et al., 2018; Mimica et al., 2023), and significant scores above zero confirm that our automated pipeline is able to detect semantic tuning without manual data curation.

**Figure 3:**
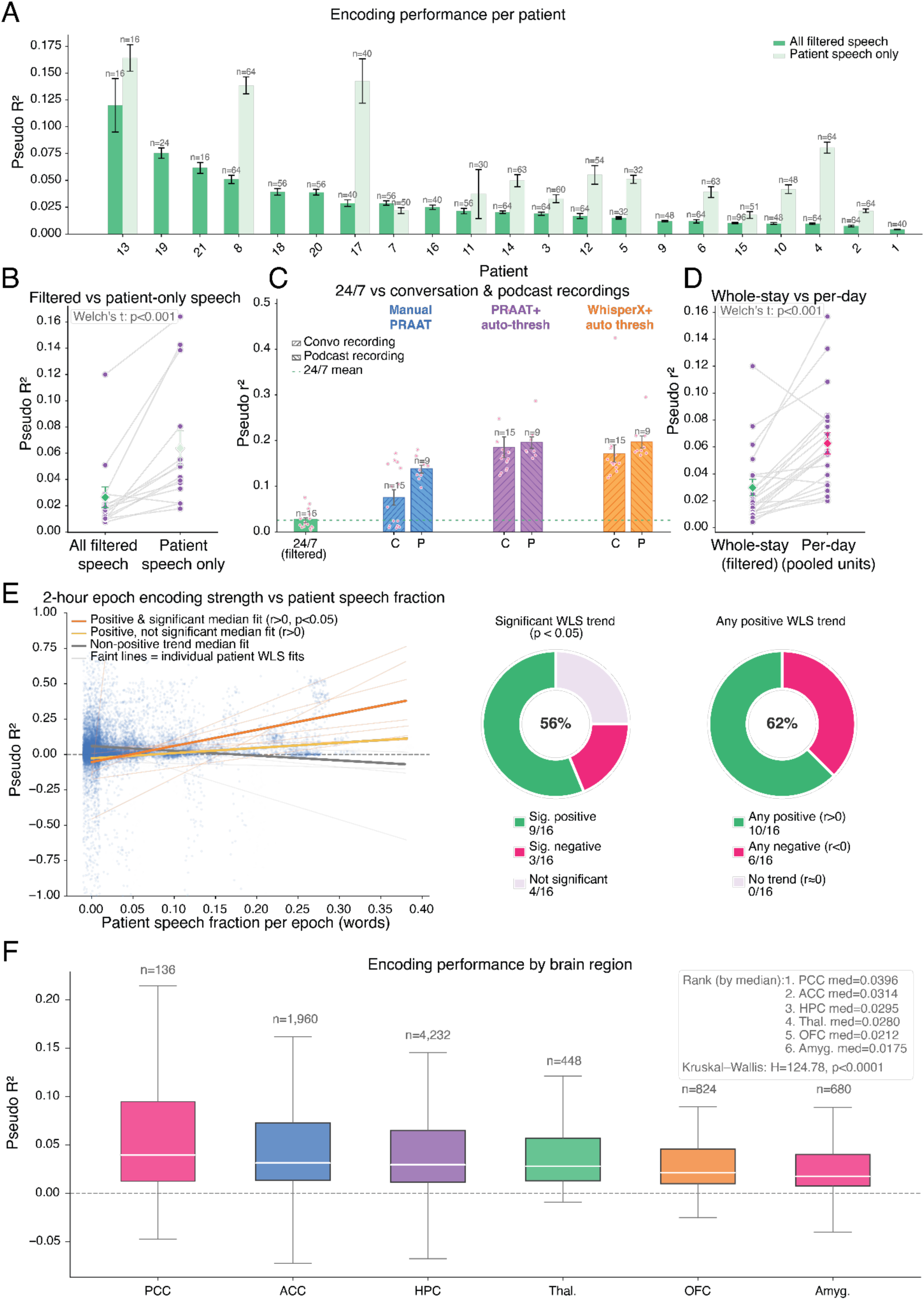
Significant Semantic Coding from Naturalistic Speech Data. **A.** Bar plot showing average semantic encoding performance per patient across all significant units with standard error bars, sorted by model performance. Number of significant units are listed above each bar. **B.** Paired line plot comparing semantic encoding performance between all speech and patient-only speech conditions. **C.** Grouped bar plot showing comparative semantic encoding performance between the naturalistic speech and experimental datasets (conversation + podcast), with automated conditions for each task. **D.** Paired line plot comparing semantic encoding performance between all speech and within-day speech conditions. **E.** Line plot with best fit lines and summary charts showing relationship between epoch encoding strength and portion of time patient spent speaking. **F.** Boxplot showing semantic encoding model performance distribution by brain region.

We next compared encoding performance for patient-only speech identified by our video diarization pipeline with all ambient speech. We overall found encoding performance to be stronger for produced speech than all speech (**Figure 3B**). Specifically, we see a 2.42-fold increase (from 0.026 to 0.064) in pseudo R^2^ scores for patient-spoken speech. This finding is especially notable considering our patient-only models were trained on 50.52 (+/ 12.30) times less data on average. One reason produced speech may be more strongly encoded is that it requires attention, while ambient speech may be ignored (Mesgarani & Chang, 2012) .

### Comparison with controlled tasks

We next compared embeddings derived from incidental speech to those derived from two highly controlled tasks: (1) an unconstrained conversation task (Chavez et al., 2025; Yan et al., 2026), and (2) a podcast listening task (Franch et al., 2026; Katlowitz et al., 2026, **Methods**). We found that encoding strength is significantly higher for our controlled datasets, with a 4.43 (± 1.67)-fold average greater encoding strength for the conversation task and a 8.59 (± 3.12)-fold greater encoding strength for the podcast task (**Figure 3C**).

To gain insight into why the controlled tasks produce stronger encoding, we replicated this analysis using a version of both controlled datasets with autothresholded spikes instead of manually curated spikes. We found that using autothresholded spikes significantly improves encoder performance, with a 6.62 (± 1.58)-fold average increase for the conversation task and a 1.47 (± 0.12)-fold increase for the podcast task. Prior work has demonstrated that multiunit activity performs comparably to sorted spikes for population coding tasks and may even filter out useful signal (Kloosterman et al., 2014; Trautmann et al., 2019), which may explain why spike sorting does not improve our model performance in the aggregate.

We then tested a version using uncorrected WhisperX transcripts rather than hand-labelled transcripts. We observed no significant difference in performance from using uncorrected WhisperX transcripts compared to hand-labelled ones, with 0.95 (± 0.037)-fold change in the conversation task and 1.01 (± 0.01)-fold change in the podcast task. This result suggests manual annotation may have modest, if any, benefits for derivation of semantic encoding models. Together, these results suggest that poorer performance in the naturalistic dataset likely occurs for reasons other than the limitations of automated data processing–such as representational drift or patient attention variability.

### Variability in encoding over time

We next evaluated how pooling data across days affects our model performance compared to training models on single-day data. We trained encoding models from each individual day in each patient’s stay and compared them to models trained on the entire patient stay (averaging 6.72 days, ranging from 6-11 days). We find that models trained on individual days performed significantly better than models trained with data from all days, with a 3.06 (± 0.44)-fold greater pseudo R^2^ scores on average (**Figure 3D**). Thus, day-to-day variability is a major driver of variation in encoding model performance.

While during formal experiments we are able to limit the number of distractions and ensure patients are focused on the task, directing patient attention is not possible in a naturalistic setting, and neural tuning to semantics may differ based on the stimulus the patient is (or is not) paying attention to (Mesgarani & Chang, 2012). To probe attentional changes in model performance, we trained encoding models on two-hour epochs across the patient stay, such that we could compare encoding strength at 2 PM and 7 PM on the same day, for example. We assessed whether epoch encoding strength correlated with the portion of time in the epoch the patient spent speaking, which can potentially be viewed as a proxy for patient attention and engagement with ambient speech. For example, the portion would be 0 when the patient is passively listening to the television while engaged in a separate activity, while it could be as high as 0.7 when they are actively engaged in conversation with another person. We find that, when regressing patient speech fraction against epoch encoding strength via weighted least squares (see: **Methods**), 62% of patients have a positive trend (56% significant), while 38% of patients have a negative trend (19% significant, **Figure 3E**). While this result may be a consequence of our units being significantly more tuned to patient speech, it may also indicate that overall, semantic tuning in our units is significantly correlated with the portion of patient speech engagement, potentially indicating that patient engagement and/or attention is a significant factor in developing high-performing semantic coding models.

### Comparing Brain Regions

We found all recorded brain regions (median pseudo R2 > 0) generate significant semantic encoding models from the naturalistic speech dataset. We found PCC, ACC and HPC to be the highest performing brain regions, with the remaining brain regions ranked in the following decreasing order: Thalamus, OFC, and amygdala (**Figure 3F**). This result corroborates findings from existing literature implicating hippocampus in higher-level semantic/symbolic language processing and findings of distributed semantic networks throughout the brain (e.g., Huth et al., 2016).

### Significant semantic decoding in the continuous speech dataset

Decoding models of semantics play a complementary role to encoding models and are especially interesting for BCI purposes (Willett et al., 2023; Silva et al., 2024; Goldstein et al., 2025; Chen et al., 2024). To simplify the semantic decoding problem, we classified all spoken words into one of ten semantic categories (**Methods**; cf. Franch et al., 2026). We evaluated decoder model performance using overall accuracy scores (ratio of correctly identified words to total number of words). With ten semantic categories, chance performance was 10%.

Our pipeline produces significant decoding. Models for all patients but one decode semantics with more than 20% accuracy (average accuracy: 20.9%. ± 0.08%, **Figure 4A**). In contrast to our encoding models, we find that decoding is more accurate (although modestly so) for all speech versus patient-only speech. Specifically, we see 18% (± 2.6%) lower classification accuracy for patient-only speech (**Figure 4B**).

**Figure 4:**
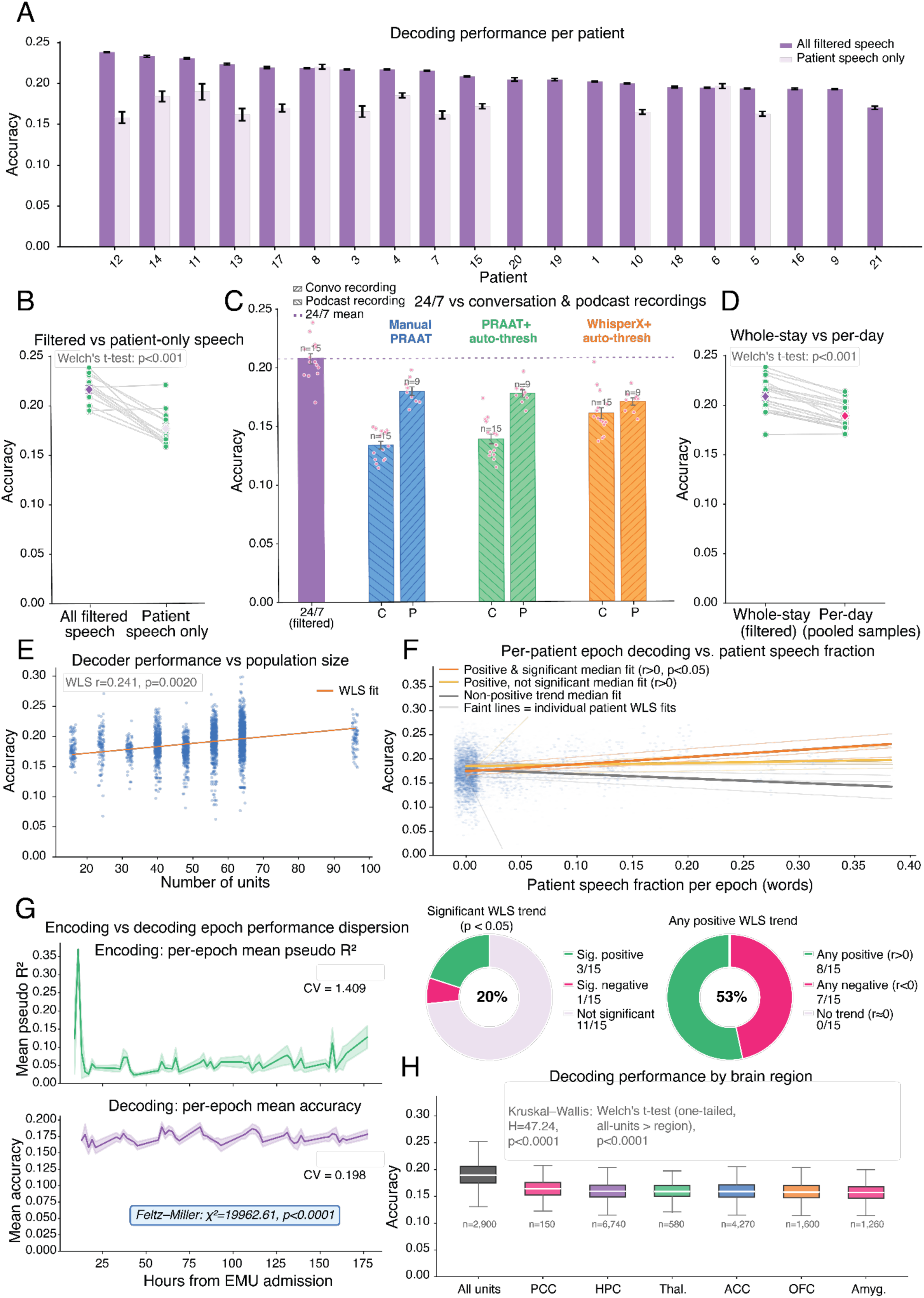
Significant Semantic Decoding from Naturalistic Speech Data. **A.** Bar plot showing average semantic decoding performance per patient across all significant units with standard error bars, sorted by model performance. Number of significant models are listed above each bar. **B.** Paired line plot comparing semantic decoding performance between all speech and patient-only speech conditions. **C.** Grouped bar plot showing comparative semantic decoding performance between the naturalistic speech and experimental datasets (conversation + podcast), with automated conditions for each task. **D.** Paired line plot comparing semantic decoding performance between all speech and within-day speech conditions. **E.** Scatterplot with line of best fit highlighting trend between unit population size and model performance. **F.** Line plot with best fit lines and summary charts showing relationship between epoch decoding strength and portion of time patient spent speaking. **G.** Line plots showcasing epoch model performance variation over time. Boxplot showing semantic decoding model performance distribution by brain region. **H.** Boxplot showing semantic decoding model performance distribution by brain region.

### Decoding: comparison with ground-truth datasets

Unlike encoding, decoding models trained on all speech perform better than models trained in our ground-truth datasets: we saw a 36% (± 1.8%) lower performance for the conversation task and an 11% (± 2.5%) lower performance for the podcast task (**Figure 4C**). Decoding may be more robust to confounds (patient attention, variable neural quality, etc.) not found in our controlled tasks than our single-unit confounds are. As a result, population decoding models may benefit more from the additional training data available via automated parsing from naturalistic environments.

As with our encoding analyses, manual curation of spikes and transcript labels do not significantly improve the accuracy of semantic decoder models for our controlled tasks. We find no significant difference (i.e. overlapping standard errors) in the decoder accuracy for models trained with sorted versus autothresholded spikes. Moreover, while we find decoder performance is 4.0% (± 1.3%) less accurate with automated transcripts in the podcast task, decoder performance is actually slightly more accurate (18.3% ± 3.2%) with automated transcripts in the conversation task. Overall, decoders appear to be more robust to the inaccuracies produced by automated neural and behavioral processing models than encoding models.

### Decoding: variability in performance over time

Decoding models are more stable over time than encoding models. Indeed, single-day model performance is 9% (3.5%) worse than full multi-day dataset decoding (**Figure 4D**). Thus, the benefit of additional training data outweighs the impediment of cross-day neural variability.

We also find that decoding accuracy is better for patients with more recorded units than fewer (**Figure 4E**). This result provides further support for the notion that larger populations of neural data allow for the training of more accurate models of human behavior that take advantage of the redundancy of single-neuron tuning (Stevenson & Kording, 2011; Stringer et al., 2019; Trautmann et al., 2019) and are robust to failures and fluctuations that occur at the single-unit level.

We also assessed the variability of semantic decoding across the two-hour epoch scale and probed whether the same attentional variation as in encoding emerged. Overall, we find a similar if less pronounced trend, where 53% of patients have a positive trend (20% significant), whilst 47% of patients have a negative trend (6% significant, **Figure 4F**). This result mirrors our encoding findings and corroborates the idea that patient engagement may improve the performance of our semantic coding models. Decoding performance is also more stable across 2-hour epochs than encoding performance: our epoch decoding performance has an 85.9% (±.16%) smaller coefficient of variation than our epoch encoding performance (p < 0.0001; Feltz-Miller test, see: **Methods**, **Figure 4G**).

### Decoding: comparing Brain Regions

We trained decoding models separately for each brain region. All brain regions performed significantly above chance, but worse than models trained using all available units across regions (p<0.001). As with the encoding models (see above), the PCC and HPC are the highest performing regions, followed by the thalamus and ACC (**Figure 4H**).

### Encoding model performance as a function of dataset size

The scale of our dataset lets us determine the effect of data quantity on performance. When we randomly decimate our per-day datasets to portions ranging from 1% to 90% of the data (**Methods**), we generally find that additional training data improves encoding model performance (**Figure 5A**). That performance shows marginal decline, although we do not detect any saturation. Our best fit curve is modelled after the Hill equation, which is commonly used to sigmoidal biological relationships (**Methods**). Moreover, we found the Hill equation to significantly fit our data according to permutation testing (p<0.01) and also better explained experimental data than a weighted least squares fit (Loss_hill < Loss_wls; dLoss==90.4).

**Figure 5:**
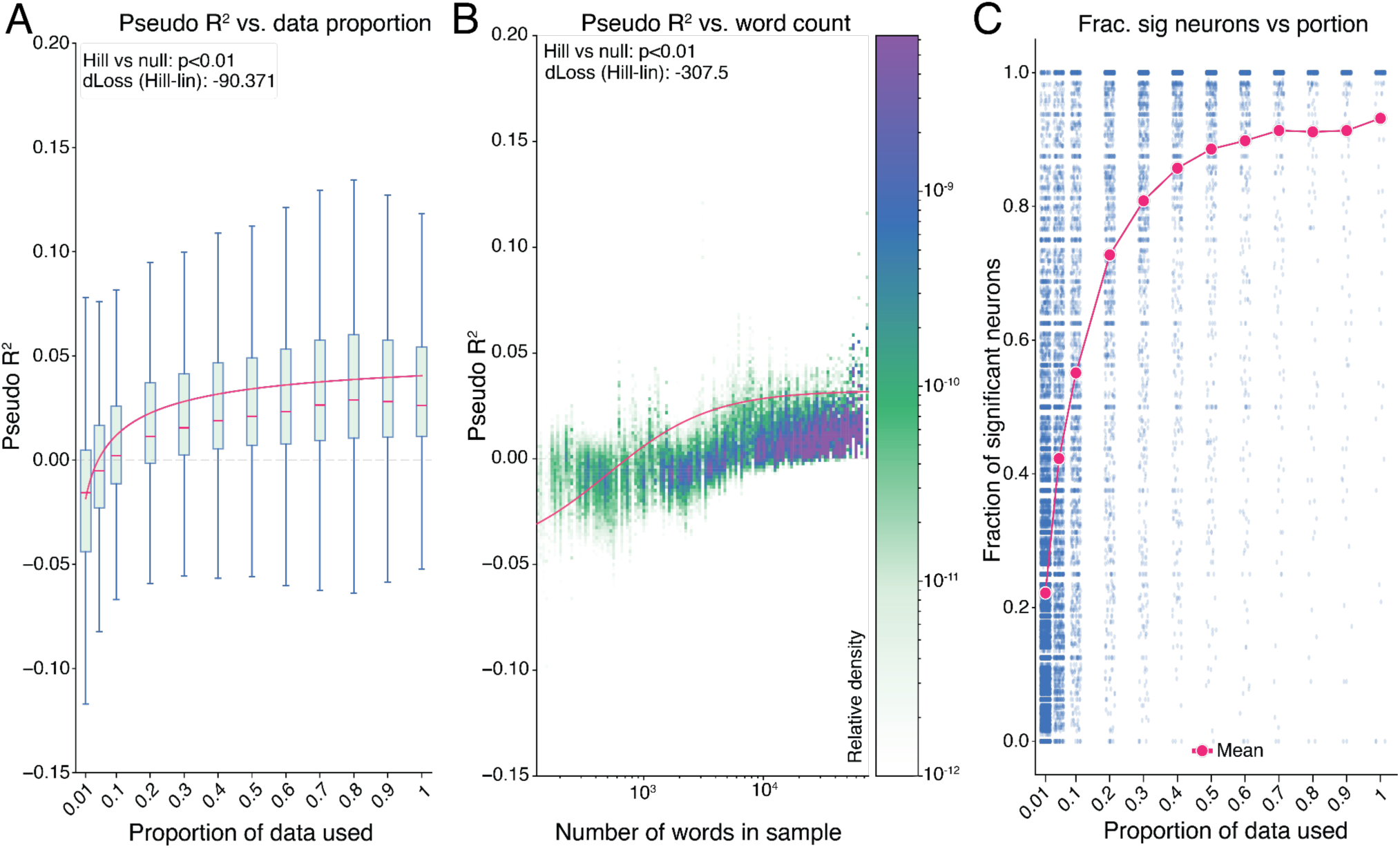
Decimation Analysis. **A.** Boxplot with Hill curve showing semantic encoding performance growth as function of dataset portion used for training. **B.** 2D Histogram with Hill curve showing semantic encoding performance growth as function of words in sample. **C.** Scatterplot of fraction of significant semantic encoding units as a function of dataset portions, with portion averages as an overlaid line plot.

We next quantified the minimum amount of data needed to generate statistically meaningful results. We calculate pseudo R^2^ by comparing it to a simpler model using word duration only. We find that model performance exceeds durational null performance when using at least 4.7% (± 0.04%) of the dataset on average.

We then repeated these analyses using number of words, instead of number of minutes. As with minutes, we find a clear logistic growth curve, with performance exceeding null at around 724 (± 3) words (**Figure 5B**). The Hill equation similarly significantly fit our data and performed better than a simple linear fit (Loss_hill < Loss_wls; dLoss=-307.5).

We find that the proportion of units significantly tuned to our semantic embeddings also increased according to a logistic growth curve, with 1% of data yielding an average of 22.2% (± 0.2%) significant units before saturating to an average of 91.6% (± 0.6%) of units at around 70% of each dataset (**Figure 5C**). Contextual transformer embeddings capture many aspects of language relevant to word meaning–including syntax, grammar, word order, surrounding context–within a distributed high-dimensional representation (Tenney et al., 2019), which means a unit significantly tuned to these embeddings could be responsive to any combination of these features. Moreover, the general mixed selectivity of neurons in higher cortical regions (Kobak et al., 2014) may explain how weak correlations with some dimensions of the transformer embedding space may reveal themselves at sufficient sample size. Our analysis showcases a classic power curve, in that only the most prominent effects are recognizable at smaller sample sizes whereas weaker effects only emerge when there is enough data to detect them.

### Assessing differences in coding models across time (functional drift)

We wanted to assess the stability of our encoding and decoding model performance due to known phenomena of representational drift and recording inconsistencies (Jude et al., 2022; Sussillo et al., 2016). Importantly, we wanted to address the rate of model degradation over time and whether it differed for unit encoding versus population decoding models, as such questions hold significant ramifications for the development of generalizable neural coding models of behavior.

We find that, for encoding models, model performance decreases as a function of the temporal separation between the original training day and the subsequent test day (p < 0.001, **Figure 6A**). This effect is strongest when comparing within-day testing to the next-day (+1), where the median pseudo R^2^ score dips from 0.034 (± .006) to −0.027 (± .008) and tends to grow more negative over time (R^2^ scores are normalized in reference to null performance, so decreases below zero are meaningful). Moreover, the portion of significant units plummets from 97.8% (± 1.2%) on average to 7.8% (2.4%) after just one day of separation (**Figure 6B**). Overall, we find that significant model degradation occurs for our single-unit encoding models after even just one day of temporal separation, rendering most models unreliable for downstream use cases after their original training date.

**Figure 6:**
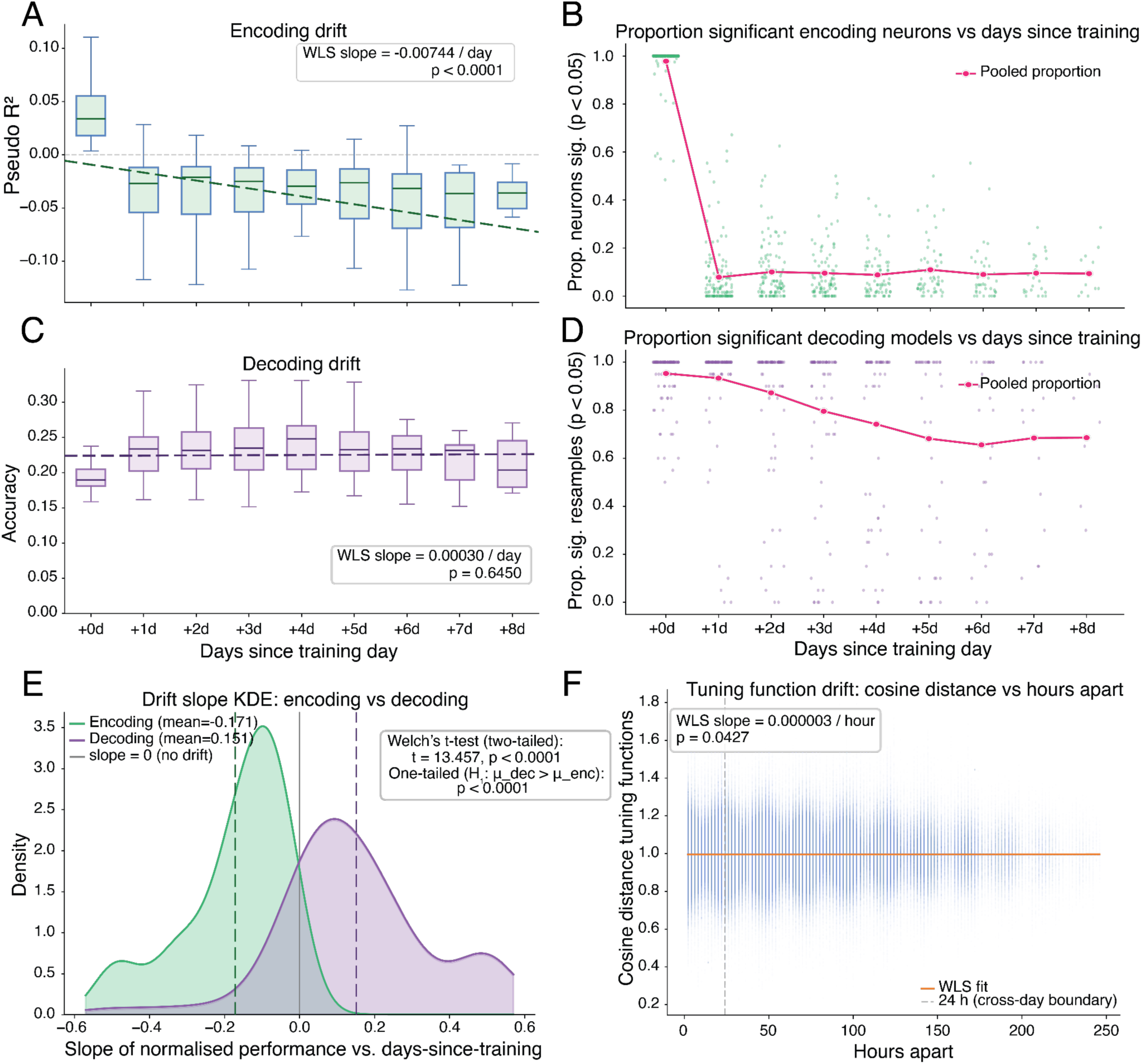
Functional Drift. **A.** Boxplot with line of best fit showing encoding model performance changes across time in days. **B.** Scatterplot of fraction of significant semantic encoding units as a function of test distance in days. **C.** Boxplot with line of best fit showing decoding model performance changes across time in days. **D.** Scatterplot of fraction of significant semantic decoding models as a function of test distance in days. **E.** Density plots showing distribution of encoding model performance slopes (green) and decoding model performance slopes (purple). **F.** Scatterplot with line of best fit showing changes in tuning function distance across time in hours.

Unlike encoding models, decoding model performance does not significantly decrease with distance in time (p= 0.6450, **Figure 6C**). However, similar to encoding, the proportion of significant models does decrease significantly over time (weighted least squares, p<0.001), albeit at a much more steady rate (**Figure 6D**). Specifically, the percent of significant models starts at 95.3% (± 1.9%) and degrades to 65.6% (± 9.1%) of models still performing with statistical significance after eight days from their original training date. The relative stability of decoding drift is apparent when we compare the distributions of regression slopes of individual model performance between encoding and decoding. We found model decoding slopes to be significantly higher than model encoding slopes (One-tailed t-test, p<0.001 **Figure 6E**).

Moreover, the overwhelming majority of encoding models have negative slopes (meaning their performance decreases over time), while the average decoding model has a positive slope.

We next investigated the differences between the tuning functions for our single-unit encoding models. The vectors of weights of our models can be thought of as tuning functions that map behavioral representations to neural activity (Chavez et al., 2025; Kriegeskorte & Douglas, 2017; Weber & Pillow, 2017). We find that as the amount of time between training sets increases, the cosine distance of our tuning functions increases very gradually (weighted least squares, p=.0427, **Figure 6F**). Thus, beyond performance loss, the temporal degradation in single-unit semantic encoding is accompanied by slight drift in the semantic tuning functions themselves. Therefore, while maximizing recording stability is an important avenue for the development of consistent semantic coding models, additional model-level corrections (such as aligning latent neural embeddings, Dabagia et al., 2022) are also likely needed to account for inherent neural variability across time.

## DISCUSSION

We developed a pipeline for processing of incidental speech data recorded 24/7 in the EMU. It records all speech and brain activity from implanted Behnke-Fried microwire bundles during the patient’s stay (6+ days). Our pipeline includes automated spike sorting, speech transcription, and video-assisted diarization. We used this pipeline to generate reliable encoding and decoding models, whose performance we quantified by using ground-truth datasets consisting of manually annotated speech and manually sorted spikes. One of the most difficult practical, if not technical, problems with such pipelines is developing tools to assess their quality. Here we used a painstaking manual curation of naturalistic data involving both hand-sorting of spikes and precise manual alignment of all speech during typical sessions. This dataset allowed us to precisely quantify the accuracy of our much larger naturalistic dataset.

We found that both encoding and decoding models are robust and reliable, and decoding models trained on longitudinal incidental speech performed better than their controlled-task counterparts. These results indicate that the level of neural responding to language is such, and of such a type, that we can infer reliable, generalizable models from passively recorded speech.

Moreover, they tell us that currently available methods are able to solve historically difficult problems in transcription, diarization, and spike sorting sufficiently well as to generate reliable models. This is important for applications in which bespoke data collection is impractical, such as in the continuous updating of brain-computer interfaces, or in scientific projects requiring large amounts of data, or long periods of time, or very infrequent words.

We found our encoding model performance scales with the size of the training dataset. Notably, even with large datasets, we did not find a saturation point for performance as a function of training set size. This result argues for the potential value of (large) naturalistic datasets in training models, especially when current approaches struggle to reach decoding accuracies useful for clinical application (Défossez et al. 2023; Tang et al., 2023; Metzger et al., 2023; Silva et al., 2024; Willett et al., 2023; Moses et al., 2023). Critically, this performance increase is achievable using automated pipelines, making the use of large datasets feasible; this is important because, as training sets grow larger, data preprocessing increasingly becomes a bottleneck.

Current approaches to modelling speech are often split between those that use single units and those that use local field potentials (LFPs). These two measures offer complementary, and partially overlapping, views of brain activity. Here we focus on spiking activity. While LFPs are relatively straightforward to analyze, and require little preprocessing, spiking activity requires detection (requiring distinguishing spikes from spike-like noise) and isolation (assigning spikes to the neurons that generate them), which is more difficult. Many automated and partially automated systems exist (Chaure et al., 2018; Chung et al., 2017; Pachitariu et al., 2024; Bucccino et al., 2020), but none approach the accuracy of careful manual annotation. Here, we show that perfect neuron-level spike isolation (as defined by that achievable by manual sorting) is not necessarily required for neural modeling, and current automated systems–specifically, spike autothresholding–are sufficient to generate reliable spike-based encoding and decoding models.

We found that we can recover significant language representations in all recorded brain regions, including the anterior cingulate cortex, hippocampus, and the centromedial head of the thalamus. These findings are inconsistent with the idea that language is strictly localized to specific language network regions, but is consistent with a growing body of neuroimaging work showing widespread brain representation of language (Huth et al., 2016; Tang & Huth, 2025; Ryskina et al., 2025; Popham et al., 2021; Caucheteux & King, 2022; Pereira et al., 2018). Our results therefore extend these ideas from neuroimaging to the domain of spiking activity. They also open the door to large data-driven studies of subtler differences between brain regions, including variation in semantic content and grammatical role.

Why did the large increase in longitudinal training data improve our population decoding models but not our channel-level encoding models? Our encoding models were fit to threshold-crossing activity from individual recording channels, rather than to spike-sorted single neurons; therefore, longitudinal nonstationarity could affect them in at least two ways. First, physical changes at the electrode–tissue interface can alter which neuron or mixture of neurons contributes to threshold crossings on a given channel. Thus, the same channel across days may not correspond to the same local neural source, making the apparent channel–stimulus relationship unstable even without any change in the underlying neural representation (Perge et al., 2013). Second, even in the absence of physical electrode drift, the neural representation itself may change over time. Such representational drift has been observed across sensorimotor and sensory systems (Rule et al., 2019; Schoonover et al., 2021; Deitch et al., 2021). This is important because an encoding model asks what a particular channel’s threshold-crossing activity is tuned to. Increasing the amount of training data should improve estimation by expanding the sampled stimulus space and providing more observations of similar semantic conditions, but only under the assumption that each channel’s tuning function remains sufficiently stationary. If a channel’s apparent tuning changes because the electrode samples a different local neuronal mixture, or because the same neurons undergo true representational drift, then pooling more longitudinal data can combine inconsistent channel–stimulus mappings rather than improve the model.

Population decoding models are expected to be more robust to this form of drift. Unlike single-unit encoding models, a decoder is not required to preserve a fixed relationship between a semantic feature and any one neuron. Instead, it can exploit redundancy across units, down-weight unstable or weakly informative units, and read out information from population dimensions that remain predictive over time. This interpretation is consistent with work in mouse visual cortex showing that individual neurons can change their activity rates and tuning over minutes to days, while the relationships among population activity patterns remain relatively stable and stereotyped (Deitch et al., 2021). It is also consistent with evidence that sensory cortical population codes can become reliable through redundancy and structured inter-area co-fluctuations, making population decoding robust to day-to-day variability in individual cells (Ebrahimi et al., 2022). More generally, task-relevant population dynamics can be recovered without relying on stable spike sorting of individual units (Trautmann et al., 2019), and low-dimensional latent dynamics or manifolds can remain stable even when the recorded units themselves change, supporting reliable decoding over long timescales (Gallego et al., 2020; Karpowicz et al., 2025). Longitudinal speech-neuroprosthesis work further supports this population-level view, showing that decoders trained and adapted using cross-day data can maintain high accuracy over months of use (Card et al., 2024). Thus, our results are consistent with an encoding–decoding asymmetry: single-unit encoding models directly expose drift in unit-level tuning, whereas population decoding models can average over, reweight, or align across unstable units to recover more stable population-level structure. This difference may explain why additional longitudinal data improved decoding performance but provided little benefit for naturalistic encoding models.

Variability in attention engagement is a major potential source of noise in incidental language studies. During dedicated tasks, patients attend closely; in the naturalistic speech settings, the patient decides what to attend to and when. It is highly unlikely that the patient is attending to the same degree at all times, or even attending at all most of the time. Neural activity likely covaries less with speech the patient is not attending to (Mesgarani & Chang et. al, 2012; Parra et al., 2016; Francart et al., 2019). Thus, one area for improvement in future automated models would be developing techniques for monitoring and controlling for attention levels, such as gaze-tracking or pupilometry (Gehmacher et al., 2024; **Zekveld, Koelewijn & Kramer 2018**). Future studies will need to address this shortcoming as well as others–including deficits in single unit recording stability–in order to develop models from passively-recorded naturalistic data that are not only scientifically meaningful but also clinically usable.

## METHODS

### M1. Electrophysiology Recording

All recorded epilepsy patients were undergoing neural monitoring for seizure localization at the Epilepsy Monitoring Unit (EMU) at Baylor St. Luke’s Medical Center in Houston, TX. Patients had Behnke-Fried intracranial electrodes placed in various regions of the brain that differed per patient, with a subset of depth electrodes including microwires from which single unit data was recorded and analyzed for this study. Overall general methods for this study are found in our prior published work (Provenza et al., 2024).

### M2. Voice Activity Detection

To identify sections of our recording containing human speech, we used silero vad (Silero Team, 2024), which is widely used in many speech processing pipelines and is considered state-of-the-art for voice activity detection, including in noisy, naturalistic environments.

### M3. Sentence Deduplication

Our audio streams included two room mics and the monitor stream, and also included the TV audio and a lapel mic for select patients. To prevent double counting the same speech picked up from multiple modalities, we performed the following deduplication procedure.

We indexed each sentence into an interval tree for efficient lookup of overlapping sentences. In cases where two sentences from different modalities overlapped, the sentence with the highest WhisperX quality score was maintained while the others were removed if their overlap ratio was greater than 0.5. This removes most duplicate sentences while allowing for some degree of temporal overlap between audio picked up from different recording devices. The overlap ratio of two sentences A, B is defined as:

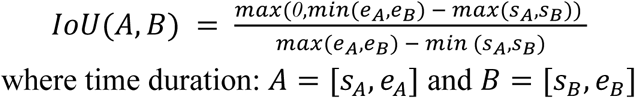

### M4. Quality Metrics

For every sentence, we calculated its corresponding CTC log-likelihood, WhisperX quality score, spectral entropy, and DNSMOS quality scores. We identified outlier sentences using the interquartile range of each distribution and removed these sentences.. Depending on the patient, this resulted in 16-65% of our sentences being considered low quality and filtered out before downstream analysis.

The connectionist temporal classification (CTC) loss quantifies the likelihood of a transcription given its associated audio segment. Using a pretrained wav2vec model, which is known to generate phonetically accurate acoustic embeddings of speech [8], we calculated embeddings of each audio segment and found its CTC loss value with its corresponding WhisperX transcript.

We specifically used the wav2vec2-large-960h-lv60-self weights for the model. The CTC loss for transcript *y* and audio segment *x* is formalized below as:

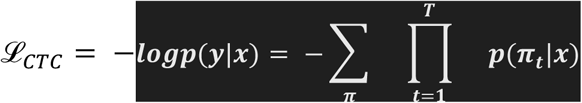

where *π*_*t*_ is a symbol set in the set of all acoustic frame-level paths that lead to our target transcript. The precise formulation of frame-level paths as well as the algorithmic implementation of the loss can be found in the original Connectionist Temporal Classification paper [9]. Our CTC log-likehood score is simply defined as the negative of the CTC loss value:

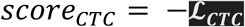

Spectral entropy measures the complexity and randomness of a signal and can serve as a proxy for assessing the noisiness of an audio segment (clean speech is typically concentrated in select frequency bands), with the aim being to exclude especially noisy segments. To compute spectral entropy, we transformed each audio segment into a power spectrogram time series with the short-time Fourier transform (STFT). We normalized each spectrogram across frequency bins to form a spectral probability distribution per time bin *P*(*f*, *t*). We then computed the Shannon entropy of this distribution per time bin:

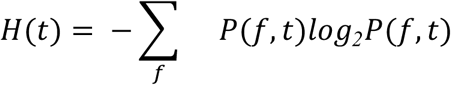

We took the average of *H*(*t*) across time bins to calculate a segment-wide measure of spectral dispersion which we report as the spectral entropy score.

We also filtered audio segments via quality scores provided directly by the WhisperX pipeline, which quantify the pipeline’s alignment confidence between audio and its associated transcript. We lastly filtered using scores provided by DNSMOS, a neural estimator of perceptual speech quality trained to approximate human-labeled quality scores on measures of speech clarity and background noise. We refer readers to the respective papers for specifics on how these scores are calculated [3, 10].

Together these filtration metrics provide a means to exclude low-quality transcripts that would negatively impact our downstream modeling performance without resorting to extensive manual checks of our dataset. We aimed to capture measures across orthogonal modes of contamination, especially those concerning raw audio quality and transcript alignment probability.

### M5. Spike Autothresholding

To capture spike counts across channels per word, we used the following bisection autothresholding algorithm:

Firstly, raw extracellular voltage traces were bandpass filtered to spike band range (300-6000 Hz) using a zero-phase forwards and backwards pass of the 4th order Butterworth filter.

For each channel we calculated the median absolute deviance (MAD) of the bandpassed voltage signal as the noise scale for calculating the spike threshold:

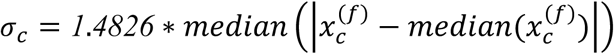

Spikes were detected as local minima in the signal below -*kσ*_3_, where *k* is solved via bisection. To prevent double-counting of spikes an absolute refractory period of 1 ms was required for the detection of a subsequent spike.

To solve *k* we calculated the global firing rate across channels for every bisection iteration of *k* and iterated until the global firing rate was within 0.5 Hz of 20 Hz or the maximum number of iterations had been exceeded (25). The global firing rate of *k* is defined as:

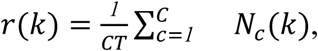

where *C* is total channels, *T* is recording duration, and *N*_3_(*k*) is the recorded number of spikes for a particular channel *c* and constant *k*.

Detected spike times were converted to a binary spike train per channel sampled at 1 kHz where the value was set to 1 for each 1 ms time bin where any number of spikes occurred.

Subsequently, spike counts per word was calculated as the number of spikes occurring during the total duration of the word.

### M6. Emotion Embeddings

To calculate emotion embeddings for all words in our dataset, we first calculated emotion embeddings for associated audio chunks. Audio segments were resampled to 16 kHz and audEERING (audeering/wav2vec2-large-robust-12-ft-emotion-msp-dim) was run for two-second long chunks with a one second stride. This resulted in a single valence, arousal, and dominance score for each audio chunk.

To get word-level valence, arousal, and dominance scores, we calculated the overlap time of the word with associated chunks:

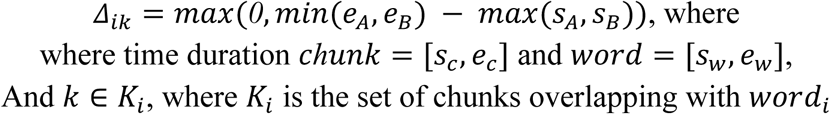

The weighted averaging equation for calculating word-level valence, arousal, and dominance scores is as below:

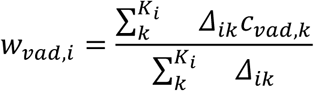

### M7. Poisson GLM

Our semantic encoding model involved using GPT-2 embeddings to predict firing rate. Our model inputs are defined below for every *i*th input:

- feature vector *x_i_* ∈ *ℜ*^8^, where *d* is the dimensionality of the principal components (PCs) of the last GPT-2 hidden state (1280 → 100). PCs were used to maximize information across dimensions and reduce model complexity.
- spike count vector *y_i_* ∈ *ℜ*^9^, where *K* is the number of channels recorded from and each value is the associated spike count for word *i*
- word duration *Δ_i_* in milliseconds

- offset *o_i_* = *log*(*Δ_i_*)

Our GLM predicts the associated spike count of an input vector with the log link function:

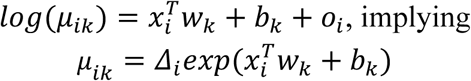

Parameters were estimated by minimizing the summed Poisson negative log-likelihood across observations and neurons with an L2 (ridge) penalty:

All *K* neurons were fit jointly using matrix operations in PyTorch, yielding a single scalar loss optimized with the L-BFGS algorithm.

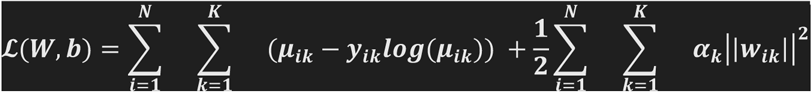

We used 5-fold outer cross-validation to evaluate held-out performance. Within each outer training split, we selected *α**_k_*** via 5-fold inner cross-validation by maximizing mean validation log-likelihood per neuron over a grid of candidate α values. To accelerate hyperparameter search, we warm-started optimization across successive α values (evaluated from high to low regularization) by initializing each fit with the solution from the previous α on the same fold. Values for α were selected from 30 candidates log-spaced in the range S*10*^+*3*^, *10^3^*T. *M8. Pseudo R^2^*

Pseudo R^2^ values were calculated as the average Pseudo R^2^ value across 5 held out test sets in a 5-fold cross validated scheme. Pseudo R^2^ definition is below:

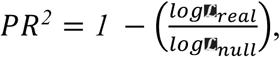

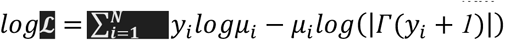, where *Γ* is the gamma function *μ_ik,real_* is the individual prediction from our Poisson GLM, whereas *μ_ik,null_* is the prediction from our null model predicting spike counts solely from average firing rate of each channel, formalized as:

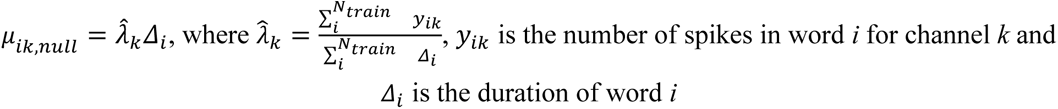

### M9. XGBoost Decoder

To decode semantic category labels from population firing rate activity, we trained multiclass gradient-boosted decision tree classifiers using XGBoost. For each word *i*, the feature vector *x_i_* is defined as the average firing rate across the word’s duration for all *K* recorded channels. Per-bin firing rate was calculated by convolving a gaussian kernel with 50 ms standard deviation across our autothresholded binary spike train. The average firing rate was calculated as the mean of all firing rates whose bins overlapped with the word duration. The output label *y_i_* is the corresponding semantic category of the word, which is classified from the word2vec embedding of the word (see: **M11**). The model was optimized using the multiclass log loss:

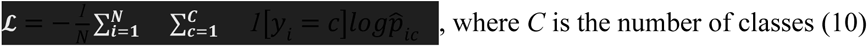

Hyperparameters including max_depth, min_child_weight, gamma, subsample, colsample_bytree, reg_lambda, reg_alpha, learning_rate, and n_estimators were tuned using 5-fold Randomized Search Cross Validation (implemented in scikit_learn). The highest performing hyperparameters were then used to train a model on the full label-stratified training set and evaluated on a single held-out test set.

To reduce prevalence bias, we excluded function words and downsampled remaining classes to the median class count via random sampling. To prevent sampling bias, we fit the XGBoost model with the above methodology for 20 resamples. To reduce compute time, hyperparameter tuning was only performed for the first 3 resamples and the remaining resamples used the hyperparameters from the initial 3 fits that maximally reduced model complexity.

### M10. F1 Macro

Performance of our semantic XGBoost decoders was evaluated using the calculation of the F1 macro score on the held-out test set:

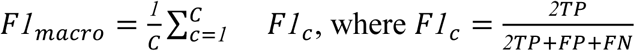

The average F1 macro along with its standard error is reported per patient for decoder comparison.

### M11. Semantic Category Classifier

Semantic categories were determined from an XGBoost classifier trained to predict class labels from a word’s word2vec embedding. Initial semantic labels were provided by the conversation dataset through clustering methods described in Chavez et al [5]. To extrapolate these labels to other datasets, we generated word2vec embeddings of each conversation word with an associated label via fastText word vectors pretrained on the English sections of Common Crawl and Wikipedia (using the model downloadable at https://dl.fbaipublicfiles.com/fasttext/vectors-crawl/cc.en.300.bin). Then, we trained an XGBoost classifier with the same randomized hyperparameter search described in **M9** on a stratified training set. The F1 macro score on the held-out test set was 0.96, indicating the semantic categories described in Chavez et al [5] are reliably recoverable from their word2vec embeddings.

This classifier was used to predict semantic category labels on the naturalistic speech and podcast datasets, which had not been included in the semantic clustering described in Chavez et al [5]. To prevent uncertain labeling from confounding downstream analysis, those words with softmax probabilities < 0.80 were left as unclassified and removed from subsequent decoding analyses.

### M12. Task Information

Two control tasks, colloquially referred to as “podcast” and “conversation”, were used to compare the relative performance gain of naturalistic speech-trained models.

The podcast task involved patients listening to a 47-minute sequence of six autobiographical stories as part of *The Moth Radio Hour*, totalling 7,346 words of natural speech. They were asked to pay full attention throughout the duration of the task session

The conversation task involved patients engaging in casual conversations with research staff. These conversations varied in time per patient but were usually between 30-60 minutes and length and covered a range of topics at the patient’s discretion. Conversations frequently included patient visitors as well if present.

### M13. Sentence Matching Algorithm

To evaluate whether a particular sentence in our corrected dataset had a match in the naturalistic speech dataset, we searched all sentences in a 60 second window around the query sentence. For all sentences with a fuzzy token set ratio >= 60 (implemented as token_set_ratio in the RapidFuzz Python package), we aligned the tokens from the corrected sentence to the naturalistic sentence to calculate the Word Error Rate (WER) between the sentences. This was done via a 2D dynamic programming algorithm that calculates the minimum amount of token substitutions (*S*), deletions (*D*), and insertions (*I*) needed to transform the reference sentence into the target sentence (token-level Levenshtein edit distance). A token was considered a match (*M*) between two sentences (i.e., edit distance of 0) based on their character-level Levenshtein distance as below for tokens *A*, *B*:

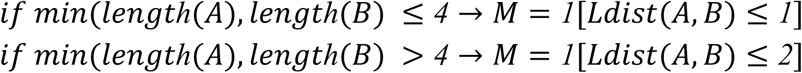

WER is formalized for a reference sentence as:

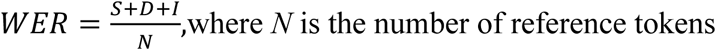

And WER similarity as:

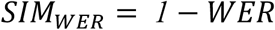

The sentence with the highest SIMwer was taken as the corrected sentences’ match. If no sentence had WER sim > 0.5 the sentence was marked unmatched first sentence found in the naturalistic dataset with *SIM_WER_* ≥ *0*.*6* was taken as the corrected sentence’s match. If there were no matches the sentence was marked unmatched.

### M14. Sentence Match Summary Statistics

The token-level Levenshtein distance described in **M12** calculates the matched words between the reference and naturalistic sentence. To define the distribution of word timestamp errors, we simply calculated the median timestamp of each word referenced to the start of its sentence and calculated the difference between the reference and naturalistic timestamp. The timestamp error for word pair *i* in sentence pair *j* is defined as:

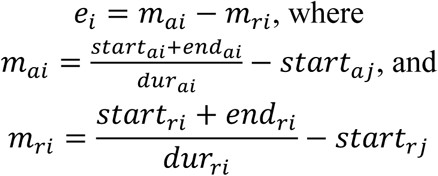

The resulting distribution describes the extent of temporal errors in sentence-level segmentation between corrected and naturalistic sentences.

We defer to the original Sentence-BERT paper for a detailed explanation of the model used to transform sentences into semantic embeddings suitable for cosine distance comparison (Reimers et al., 2019).

### M15. Permutation Testing

To evaluate whether each neuron’s GLM performance was significant compared to predictions based on word duration or other latent structural features, we randomly permuted the GPT-embeddings 50 times for each cross fold and refit our Poisson GLM with the same *α* vector. This destroyed the semantic relationship between input and output whilst maintaining word duration as labels were not permuted instead. PCA was recalculated for the newly permuted training set per fold. The p-value per fold is calculated by observing the proportion of times the log-likelihood (see: **M7**) value of our permuted model is greater than or equal to our real model:

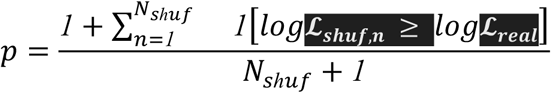

The p-value across folds for a particular neuron is calculated the same way except by using the average *log***L***L* values across folds and per permutation:

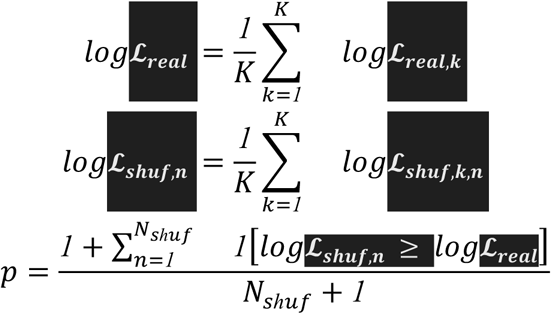

To evaluate the effect of neural preprocessing techniques on our downstream, we also sorted raw voltage traces with the algorithm included as part of the wave_clus MATLAB package (Chaure et al., 2018). The resulting sorted clusters were then manually evaluated and filtered were then manually evaluated and filtered with the wave_clus GUI. Non-spike noise was removed and each signal was classified as multi or single unit using several criteria: consistent spike waveforms, waveform shape, exponentially decaying ISI histogram, and refractory period (1 ms) violations below 5%. Sorted signals were included for both single and multi unit activity.

### M17. Decimation Analysis

To evaluate the performance growth of our encoding models, we fit Poisson GLMs from **M7** using randomly sampled proportions of the naturalistic dataset for each patient. We fit models using 1%, 5%, 10%, 20%, 30%, 40%, 50%, 60%, 70%, 80%, 90%, and 100% of the data with 100, 20, 10, 8, 7, 6, 5, 4, 3, 2, 2, and 1 resamples, respectively.

### M18. Hill Curve Fitting

To evaluate whether our performance growth for encoding models followed a logistic growth curve, we fit a predictive model of pseudo R2 as a function of data proportion/number of words used to train the model using the Hill equation:

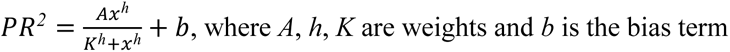

The model was fit using L-BFGS with a weighted mean squared error loss (so as to equally consider proportions regardless of sampling frequency) defined below:

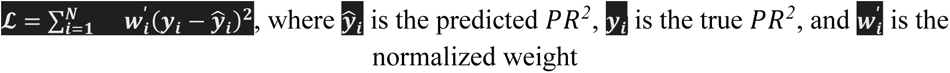

The weight term is defined as below:

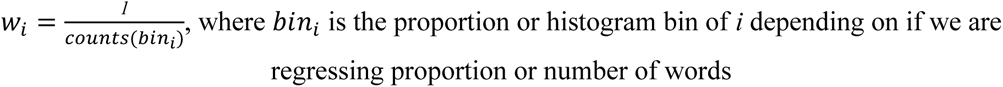

We then normalize the weight term for model fitting as below:

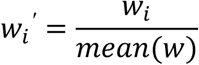

We then use the same permutation testing scheme as **M15** to evaluate the significance of our model compared to chance. We also assessed the relative performance of the Hill model to a simple linear equation with a single weight and bias by calculating the differences in their loss value, negative values indicating our Hill curve achieved a better fit than a linear equation:

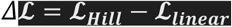

### M19. Daily Representational Drift

To assess representational drift of our semantic encoding model, we fit independent Poisson GLMs for each day of the patient’s stay according to the methodology described in **M7**. Each daily model was fit using words from 9 AM to 11 PM in local time of their respective day to coarsely separate days by patients’ general sleep-awake cycles.

To quantify representational drift, we compared the linear vector of weights for each model per-neuron using the cosine similarity score:

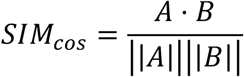

We specifically calculated the cosine similarity for all combinations of models that were trained on the same day, one day apart, two days apart, three days apart, and four days apart. To compare models trained on the same day, we trained additional models per day using only every even or odd word uttered during the day.

### M20. Embedding Extraction

To compare the relative strength of GPT-2 in training semantic encoding models, we also trained models following the methods in **M7** using embeddings derived from Sentence-BERT, Gemma 3 and word2vec. Gemma embeddings were calculated identically to GPT-2 embeddings with a 50-word context window using the gemma-3-270m version of the model, word2vec embeddings were derived identically to **M11**, and SBERT embeddings were derived identically to **M14**. For gemma and GPT-2, we used a context window of up to the prior 50 words for each recording session.

### M21. Region Decoder Sampling

To assess the relative performance of brain regions in semantic category decoding, we fit population decoders using only units from specific regions. Those units that were on the border of their intended target or otherwise missed it according to visual inspection of the post-operative, coregistered CT scan in RAVE (Magnotti et al., 2020) were excluded from this analysis. To remove unit count as a confounding factor, we fit each population decoder with only 8 units. If we had more than 8 units for a particular region, we sampled 8 units 5 times and fit 10 category-sampled models for a total of 50 fits per brain region. Otherwise, we only fit 10 category-sampled models for patient regions where there were only 8 units. To increase training efficiency, we only performed full hyperparameter search on the first three sample models per region and then picked the hyperparameters that most reduced model complexity for subsequent fits.

## Funding statement

This research was supported by the McNair Foundation and by NIH R01 MH129439, U01 NS121472, NINDS Research Education Grant Programs for Residents and Fellows in Neurology, Neurosurgery, Neuropathology, and Neuroradiology (UE5), the SNS Allan Friedman RUNN Research Grant, the NLM Training Program in Biomedical Informatics & Data Science for Predoctoral & Postdoctoral Fellows, T15LM007093-33 Gordon and Mary Cain Pediatric Neurology Research Foundation

## Competing interests

S.A.S has consulting agreements with Boston Scientific, Zimmer Biomet, Koh Young, Abbott, and Neuropace. SAS is Co-founder of Motif Neurotech

## Acknowledgements

We thank Victoria Gates and Raissa Mathura for their assistance.

